# Preparation of anti-HER-2 antibody PLGA polymer nano- ultrasound contrast agent In vitro targeting experiment

**DOI:** 10.1101/619742

**Authors:** Ji Lin, Molly Stevens, John Smith

**Author notes:** Corresponding author: Address: 631 Sumter Street, Columbia, SC 29208.

## Abstract

In this report, we demonstrated a novel method to prepare a hollow nano-targeted ultrasound contrast agent carrying anti-HER-2 antibody with high molecular weight polylactic acid-glycolic acid (PLGA) as a film-forming material, and to investigate in vitro targeting and imaging effects. We utilized the camphor as porogen, PLGA nano-ultrasound contrast agent was prepared by modified double emulsion solvent evaporation method. The general characteristics were characterized by scanning electron microscope, transmission electron microscope and laser particle size analyzer. The angiography was performed by carbodiimide method. The anti-HER-2 antibody was used to prepare the PLGA-targeted nano-ultrasound contrast agent with anti-HER-2 antibody. The in-situ imaging ability was evaluated by laser confocal scanning microscopy. Results indicate that the average particle size of PLGA nano-ultrasound contrast agent was (152.00± 58.08) nm. The particles were regular spherical, uniform in size and good in dispersion. In vitro targeting experiments showed that PLGA-targeted contrast agents with anti-HER-2 antibodies were more strongly aggregated on the surface of breast cancer cells. In vitro imaging experiments showed that the PLGA-targeted nano-ultrasound contrast imaging showed a fine and uniform point-like hyperechoic echo, and no significant attenuation of the posterior echo. This study successfully produced a PLGA-targeted nano-ultrasound contrast agent with anti-HER-2 antibody, which can specifically bind to breast cancer cells with high expression of HER-2 receptor in vitro, and the imaging effect in vitro is better.

## Introduction

In recent years, high molecular weight polymers have been widely used in medical ultrasound contrast agents due to their biocompatibility, biodegradability and good film-forming properties [1], among which polylactic acid (PLA) and polylactic acid-hydroxyl groups Polylactic-co-glycolic acid (PLGA) is the most widely used. Microbubbles made of polymer as a film-forming material envelop air or perfluoropropane gas to produce strong echo scattering, which is an ideal ultrasonic contrast agent material. However, the tumor vascular endothelial space is 400-600 nm. Traditional polymer microbubbles can only be confined to blood pool imaging, which limits their ability to detect extravascular tumor cells [2-3]. In this study, PLGA was used as the material, and camphor was used as the porogen. The nano-PLGA ultrasound contrast agent was prepared by the modified double emulsion solvent evaporation method, and the nanoparticles were combined with the fluorescently labeled anti-HER-2 antibody to prepare the targeted nano-ultrasound contrast agent. To detect its in vitro targeting ability and in vitro imaging effect on breast cancer SKBr3 cells.

## Materials and Method

Main reagents: PLGA (50:50, Jinan Biotech Co., Ltd.); natural D (+)-camphor, polyvinyl alcohol PVA1788 low viscosity type (alcohol degree 87.0∼89.0 mol/mol) and mannitol (Aladdin) Reagent Shanghai Co., Ltd.; Dichloromethane (Shanghai Runjie Chemical Reagent Co., Ltd.); Perfluoropropane Gas (Shanghai Renjieling Optical Instrument Co., Ltd.); Isopropyl Alcohol (Shenzhen Pharmaceutical Group Chemical Reagent Co., Ltd.); Ultrapure Water; Fluorescein isothiocyanate (FITC)-anti-HER-2 antibody (Shanghai Yixin Biotechnology Co., Ltd.); NHS/EDC (Billing reagent, Shanghai); phosphate buffered saline (PBS) Mercury Technology); paraformaldehyde, DAPI (Sigma, USA). Main instruments: ultrasonic cell pulverizer; RE-2000 rotary evaporator; Xiangyi H-1650 high speed centrifuge; FD-1A-50 freeze dryer; field emission scanning electron microscope (S-4800, Japan); Malvern nanometer particle size Potential analyzer (ZEN3690, Germany); laser confocal microscope (Leica TCS SP5II, Germany); ion sputtering instrument (JFC-1100, Japan). Cell lines: breast cancer SKBr3 cells, MDA-MB-231 cells, purchased from the Cell Resource Center of the Shanghai Institute of Biological Sciences, Chinese Academy of Sciences.

Preparation of PLGA nanoparticles by modified double emulsion solvent evaporation method: 1 accurately weigh 12.5 mg of natural camphor, dissolve in 5 mL of dichloromethane; weigh PLGA 125 mg, also dissolve in dichloromethane; magnetically stir to fully dissolve, get Colorless transparent solution. 2 Take 1 mL of PVA solution (3%, W/V), inject the above solution to obtain a water-oil two-phase solution; perform the first emulsification with an acoustic vibrometer under ice water bath conditions (130 W, on 4 s/off 2 s, 180 s), obtaining a milky white primary emulsion. 3 The primary emulsion was injected into 20 mL of PVA solution (3%, W/V) for the second emulsification (130 W, on 4 s/off 2 s, 180 s). 4 Pour the secondary emulsion into 100 mL of isopropanol solution (5%, V/V) and spin to evaporate to evaporate the solvent and solidify the surface of the particles. 5 The mixture obtained by rotary evaporation was placed in a 15 mL centrifuge tube and centrifuged at 12 000 r/min for 10 min; the supernatant was discarded, and the white precipitate was re-dissolved in ultrapure water and centrifuged again, 3 times; A small amount of mannitol was added to the precipitate, dispersed in 1 mL of ultrapure water, and wrapped in aluminum foil and placed in a refrigerator at −20 °C for freezing. 6 It was placed in a freeze dryer for 24 h to freeze-dry, to obtain a white powder, and a vacuum was applied to inject perfluoropropane gas. 7 Keep the product sealed and protected from light in a refrigerator at 4 °C.

The morphology of PLGA hollow nanoparticles was observed by field emission scanning electron microscopy. The PLGA hollow nanoparticles were dispersed in ultrapure water. Appropriately dispersed samples were applied to the rough surface of aluminum foil to prepare dried samples and then sprayed with an ion sputter. Gold, under the scanning electron microscope, observed the particle size, morphology, surface and dispersibility. The PLGA hollow nanoparticles were dispersed in ultrapure water by a nano-particle size potentiometer, and the hydrated particle size and polydispersity index of PLGA hollow nanoparticles were measured at room temperature. It was confirmed by transmission electron microscopy whether the particles had a hollow structure: the PLGA particles were dispersed and dropped onto a copper film covered with a carbon film, and dyed with a freshly prepared phosphotungstic acid solution (1%, W/V) under a transmission electron microscope. Observe.

The PLGA nanoparticles (1 mg/mL) dispersion was incubated with the coupling activator NHS/EDC for 30 min at room temperature, centrifuged at 16 000 r/min for 10 min; the supernatant was discarded and reconstituted in PBS for 2 times. The unreacted coupling activator NHS/EDC is removed. Disperse the activated PLGA nanoparticles in 200 µL PBS, add 25 µL of 1 µg/µL FITC-anti-HER-2 antibody, mix well, incubate for 30 min, centrifuge at 16 000 r/min for 10 min; disperse the precipitate In PBS, repeat 2 times, remove free antibody, obtain targeted nano-ultrasound contrast agent, and disperse in 50 µL PBS for use.

Preparation of cell culture and slides Two breast cancer cells (SKBr3 cells, MDA-MB-231 cells) were cultured at 37 °C, 5% CO2, and fully saturated humidity, and cultured in DMEM containing 10% fetal bovine serum. For SKBr3 cells, MDA-MB-231 cells were cultured in L-15 medium containing 10% fetal bovine serum. Both cells were cultured at a density of 1 × 104 / dish in a Petri dish with a thickness of 0.17 mm, and the experiment was started 24 hours after the cells were attached.

Take SKBr3 cells and MDA-MB-231 cells one by one, add 25 µL FITC-anti-HER-2 antibody separately; take SKBr3 fine-carrying anti-HER-2 antibody PLGA polymer nano-ultrasound contrast agent preparation and its in vitro targeting Experiments · 108 · In June 2014, Volume 23, Phase 2, and MDA-MB-231 cells were each supplemented with 25 µL of targeted nano-ultrasound contrast agent. All 4 cells were incubated for 20-30 min, then washed 3 times with PBS to remove unbound antibody and targeted nano-ultrasound contrast agent. Fixation was carried out with a 4% paraformaldehyde solution and stained with DAPI. The above operations are all carried out in the dark.

In vitro imaging effect of targeting PLGA nanoparticles: accurately weigh a certain amount of PLGA hollow nanoparticle powder, prepare a dispersion of 1 mg / mL, take 5 mL of transparent plastic sample tube filled with rubber stopper, and take 5 mL at the same time Airless water was injected into the same sample tube as a blank control group. With airless water as the sound-transparent window, real-time gray-scale imaging (central frequency 22 MHz UHF probe with a mechanical index of 0.06) was performed with a Yum MyLab Twice ultrasound system.

## Results and Discussion

The prepared PLGA nanoparticle dispersion sample was dried and sprayed with gold and observed under a scanning electron microscope. The particles were spherical, the surface was smooth, the shape was regular, the dispersion was good, and no agglomeration was observed between the particles (Fig. 1). The average particle size of the particles measured by the nanoparticle size potential analyzer was (152.00 ± 58.08) nm, and the polydisperse index (PDI) was 0.221, indicating that the particle size distribution was uniform (Fig. 2, 3).

**Figure 1.**
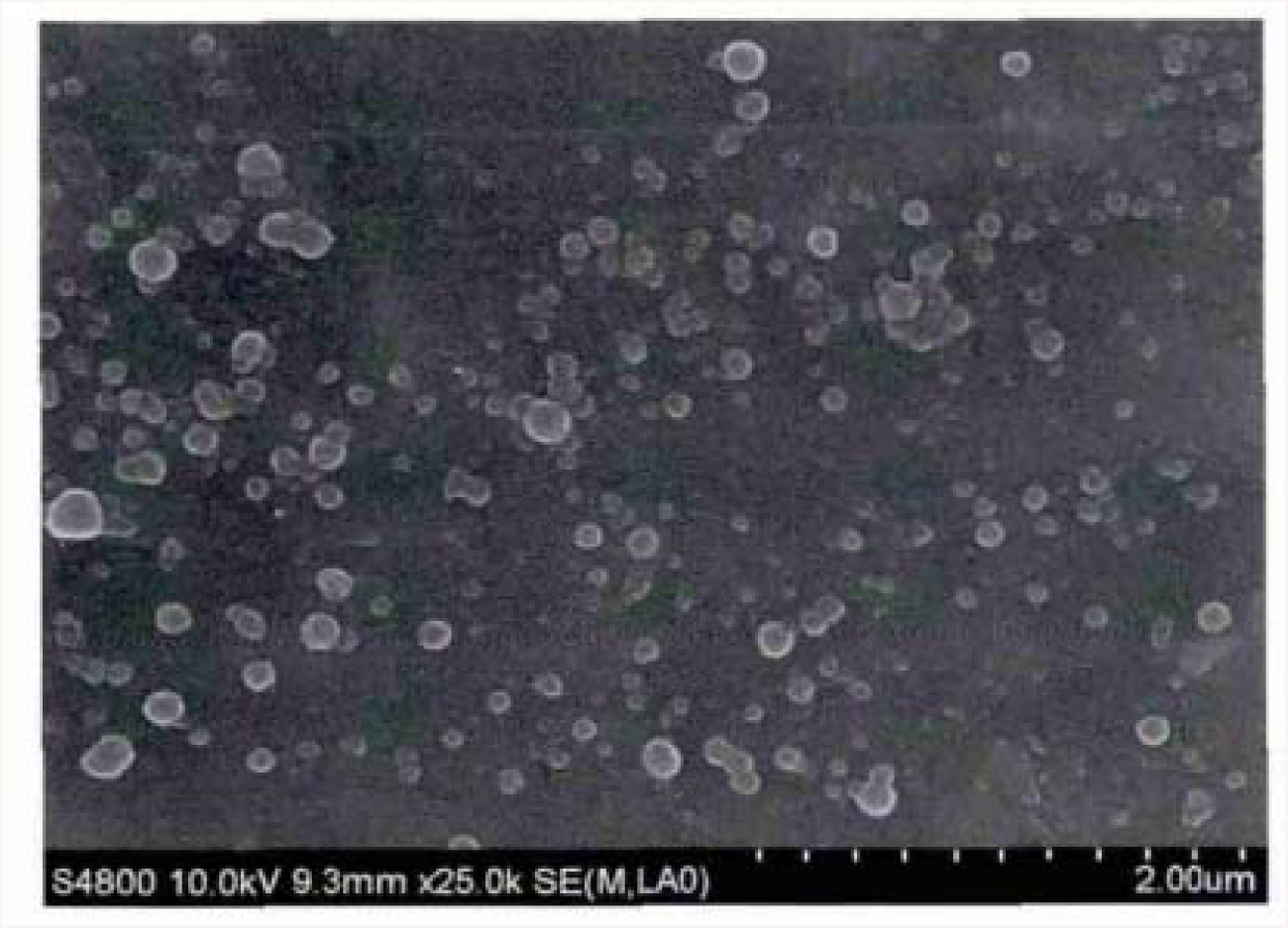
SEM Images of nanoparticles

**Figure 2.**
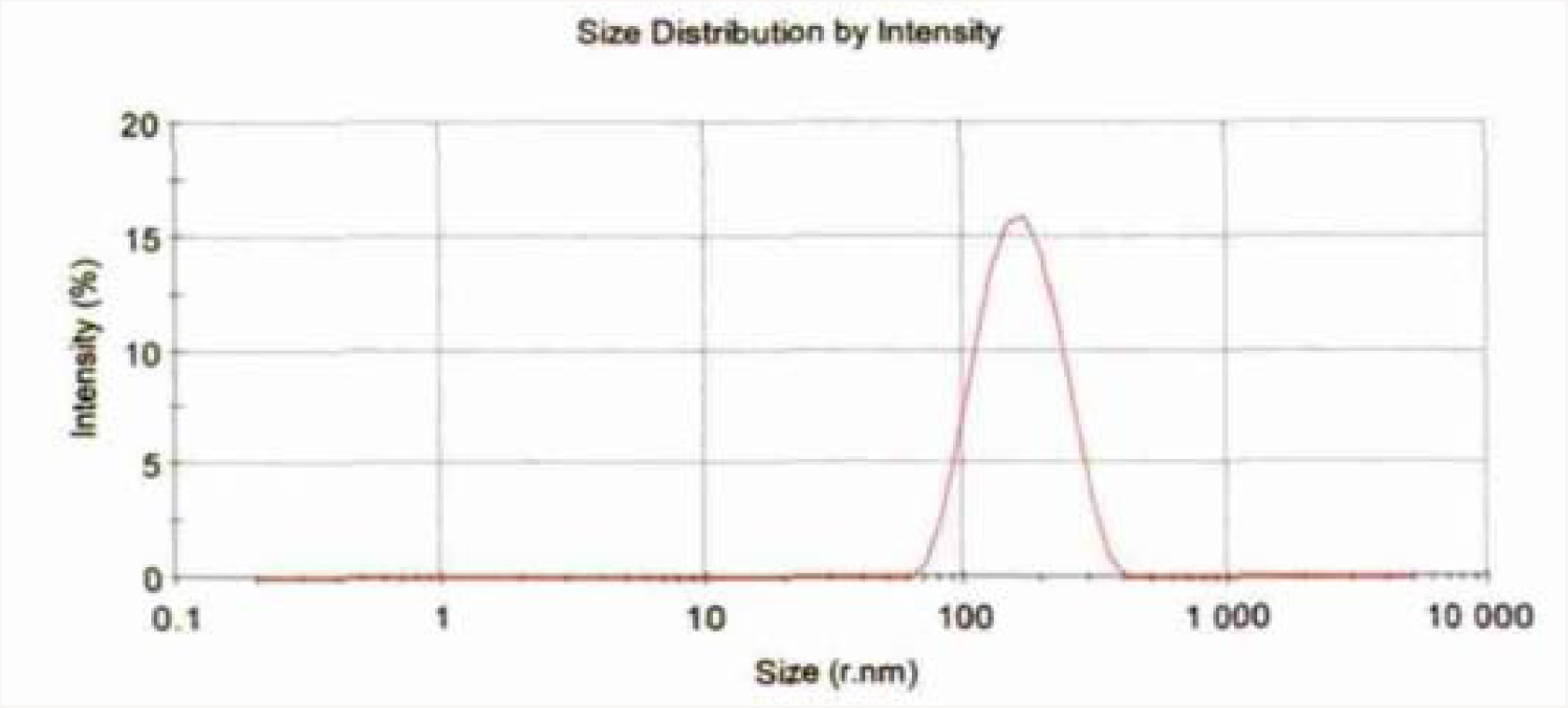
Nanoparticle size analysis by dynamic light scattering

**Figure 3.**
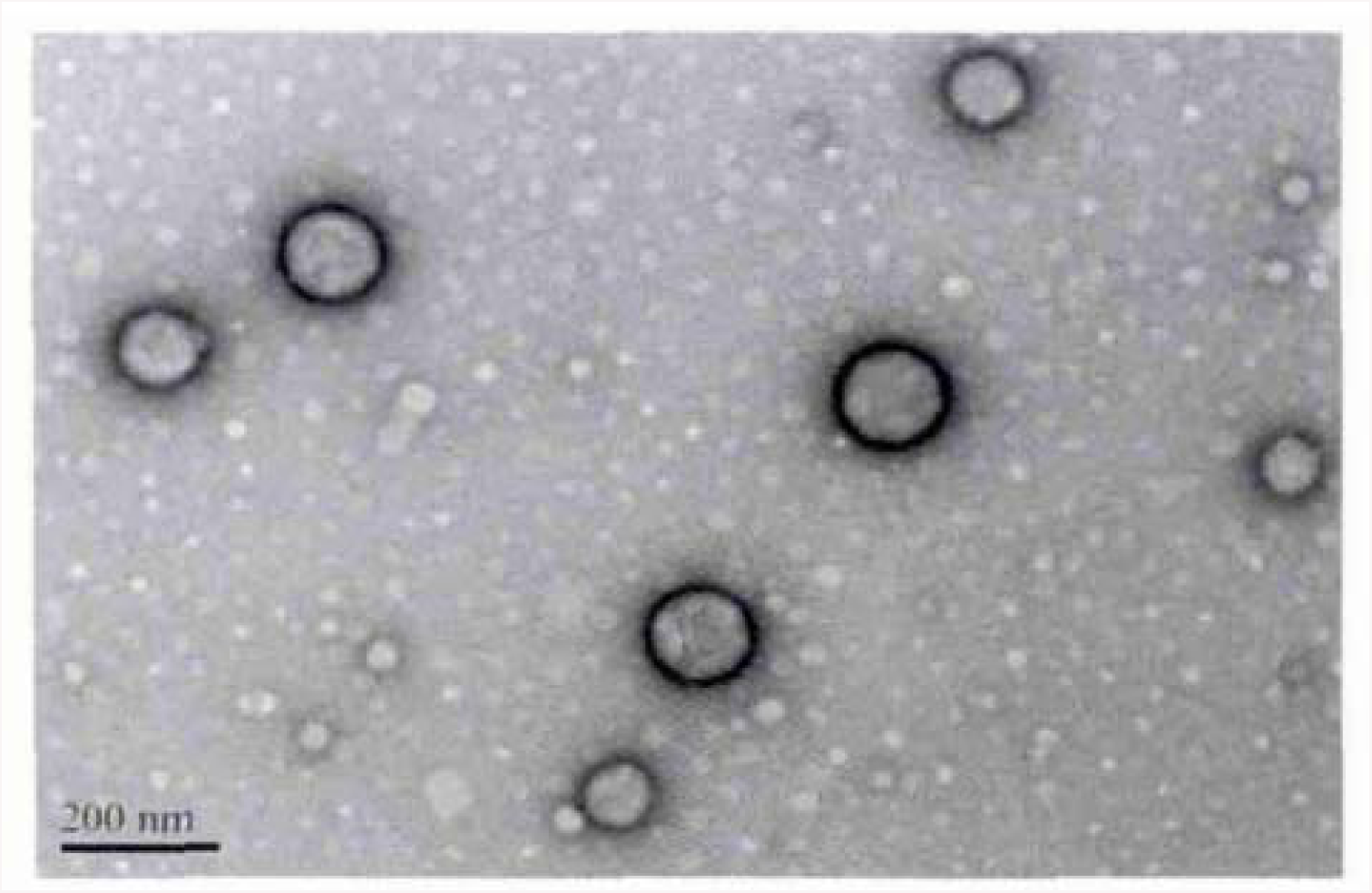
TEM images of nanoparticles loaded with fluorophore

As shown in Fig. 4A, the success rate of FITC-anti-HER-2 antibody and SKBr3 cells was observed under laser confocal microscopy. The nucleus was stimulated by DAPI staining to show blue fluorescence, and the cell membrane was excited to obtain a ring or a point. Green fluorescence, indicating that the antibody successfully binds to the cell membrane. Figure 4B shows that some of the SKBr3 cells bind to the target nano-ultrasound contrast agent, and the cell membrane shows a dot-like green fluorescence, indicating that the targeted nano-ultrasound contrast agent is more and firmly attached to the surface of breast cancer SKBr3 cells and is not eluted by PBS. In Fig. 4C and Fig. 4D, no adhesion of antibody or targeted polymer contrast agent was observed on the surface of MDA-MB-231 cell membrane, which further confirmed that the targeted contrast agent has higher expression of HER-2 receptor-expressing breast cancer cells in vitro. Strong specific affinity.

**Figure 4.**
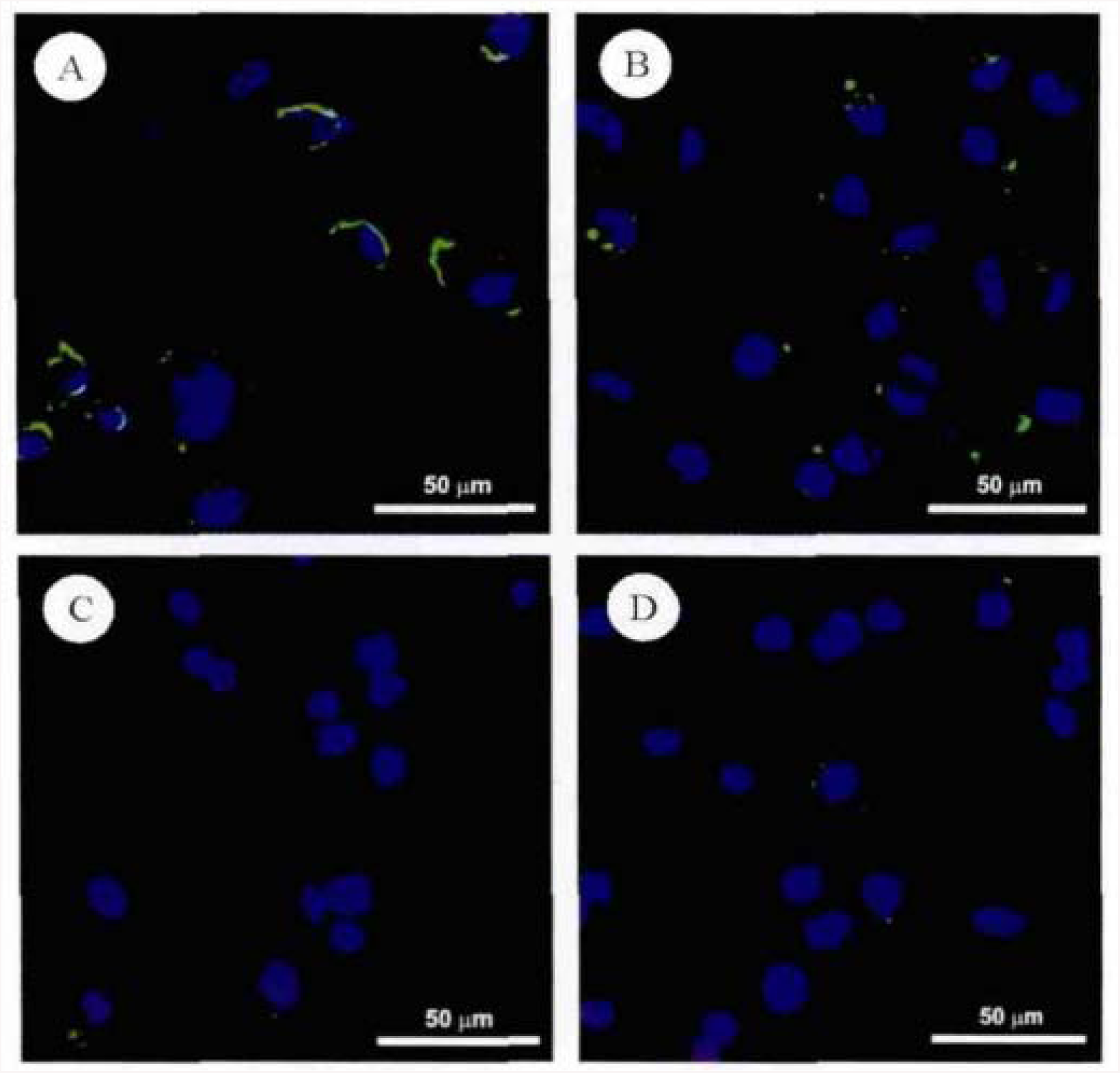
Cell uptake measured by fluorescent microscopy

The nano-targeted contrast agent with a concentration of 1 mg/mL was filled into a sample tube with a volume of 5 mL, and the sample tube containing 5 mL of degassed water was used as a reference, and the targeted nano-polymer ultrasound was observed with a high-frequency linear array probe. The ability of the contrast agent to develop in vitro. As a result, as shown in Fig. 5, the PLGA contrast agent group showed a point-like hyperechoic, which was fine and uniform, and the water of the control group showed an anechoic fluid-free area.

**Figure 5.**
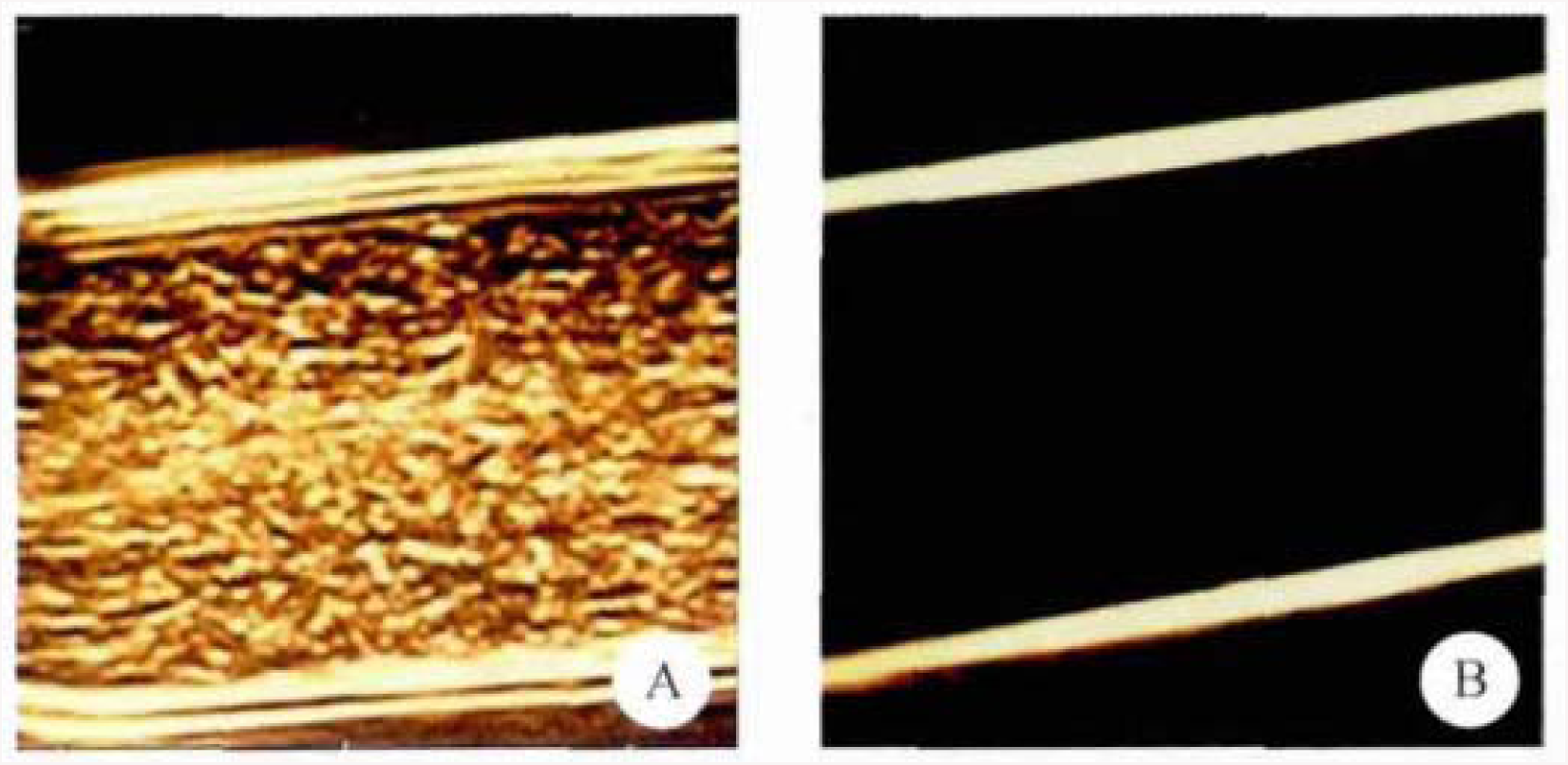
In-situ imaging of PLGA nanoparticles.

The ultrasonic contrast agent prepared by using the high-molecular polymer PLGA as a film-forming material is biocompatible and biodegradable, and can be metabolized into non-toxic carbon dioxide and water in the living body, and is harmless to the living body, and is ideal. Ultrasound contrast material has a good application prospect [4-5].

In this study, the camphor was dissolved in the primary emulsifier, and the PLGA particles coated with camphor were prepared by the modified double emulsion solvent evaporation method. After freeze-drying, the camphor sublimed and filled with perfluoropropane gas to obtain PLGA nano-ultrasound contrast agent. The surface of the particle is smooth and regular, the average particle size is (152.00± 58.08) nm, the particle size is monomodal, and the PDI is 0.221. It can theoretically penetrate the tumor vascular endothelium (400-600 nm) into the tumor tissue. It is possible to visualize extravascular target tissue through “passive targeting”, thereby overcoming the limitations of traditional microbubble contrast agents in intravascular imaging [6]. The surface of the nano-contrast prepared by PLGA contains a large amount of carboxyl groups, and the active substances such as monoclonal antibodies, ligands, and various specific short peptides are mostly proteins, which themselves contain a large amount of amino groups, and carboxyl groups and amino groups can be coupled by covalent bonds. In this study, EDC/NHS was used as a coupling activator to bind the carboxyl group on the activated PLGA to the amino group on the monoclonal antibody, and the targeted PLGA nano-sonography with anti-HER-2 antibody was successfully prepared by the carbodiimide method. And demonstrated its ability to specifically bind to breast cancer SKBr3 cells with high expression of HER-2 receptor. The nano-ultrasound contrast agent carrying the anti-HER-2 antibody is expected to be actively targeted in the body, so that the ultrasound contrast agent can be concentrated in the breast cancer tissue at a higher concentration to achieve the purpose of enhanced orientation.

Transmission electron microscopy confirmed that the nano-particles prepared in this study have a cavity structure, the presence of a cavity structure and the introduction of perfluoropropane gas can produce strong echo scattering. PLGA nano-ultrasonic agents differ greatly from conventional microbubble contrast agents in their imaging patterns, which are not suitable for imaging of traditional microbubble contrast agents on ultrasound systems. Preliminary in vitro ultrasound imaging experiments have shown that PLGA-targeted nano-contrast agents have better imaging effects under high-frequency ultrasound conditions^1-14^.

The carboxylic acid bond (—COOH) on the surface of PLGA nanoparticles and the internal cavity make it not only a targeted ultrasound contrast agent, but also a drug on the surface of the particle or a quantitative drug encapsulated inside the particle to achieve targeted tumor diagnosis and The purpose of treatment [7] lays a good foundation for the further development of PLGA nano-targeted drug-loaded contrast agents with dual functions of diagnosis and treatment.

## Conclusion

In summary, this study successfully prepared a PLGA-targeted nano-ultrasound contrast agent with anti-HER-2 antibody, which can specifically bind to breast cancer cells with high expression of HER-2 receptor in vitro, and in vitro imaging effect. better. Future efforts will focus on translating this research to the clinic.

